# Enrichment-Free Identification of Native Definitive (EnFIND) O-glycoproteome of antibodies in autoimmune diseases

**DOI:** 10.1101/2020.07.15.204511

**Authors:** Xue Sun, Jianhui Cheng, Wenmin Tian, Shuaixin Gao, Jiangtao Guo, Fanlei Hu, Hong Zhang, Xiaojun Huang, Da Yong Chen, Yang Chen, Catherine CL Wong

## Abstract

The detection of O-glycosylation at the proteome level has long been a challenging task and a roadblock for O-linked protein glycosylation research. We report an Enrichment-Free Identification of Native Definitive (EnFIND) O-glycoproteome using Trapped Ion Mobility Spectrometry coupled to TOF Mass Spectrometry (TIMS-TOF MS) for direct analysis of protein O-glycosylation in native samples with minimum sample requirement. This approach enabled separation of O-glycopeptide isomers, resolution of O-glycosites and O-glycoform, reduction of sample complexity, and increased sensitivity, thus greatly enhancing analysis of the O-glycoproteome of cell lysates, human serum and exosomes. In addition, we found that antibodies in human serum are highly O-glycosylated on variable, especially hypervariable regions and constant regions, which significantly increases antibody diversity. This method was used to successfully identify characteristic O-glycosylation features of autoimmune diseases.

## Introduction

O-glycosylation is one of the most abundant and diverse type of protein glycosylation. It mainly occurs on serine (S), threonine (T) or tyrosine (Y) residues and initiates with the addition of GlcNAc, GalNAc, Fucose, Glucose, Galactose or Mannose. The initial monosaccharide can be further elongated to form large and complex glycan structures. The O-glycosylation at specific sites contribute to the diverse regulation of protein activities and serve a variety of biological functions(Marino et al, 2010). More importantly, O-glycosylation is related to diseases including congenital disorders, cancer and autoimmune diseases(Reily et al, 2019). Analysis of O-glycosylation at the proteome level has been a challenging task for decades. O-glycopeptide ions are highly heterogeneous and their signals are suppressed in mass spectrometry by unmodified peptides due to their low abundance. Identification of O-glycopeptides by mass spectrometry is further complicated by the lack of glycan consensus structures and the existence of glycoform isomers. Substantial improvements have been made for large-scale O-glycoproteomics analysis in complex biological samples(King et al, 2017; Mao et al, 2019; Schjoldager et al, 2012; Vakhrushev et al, 2013; Yang et al, 2018; Yang et al, 2014). For example, Vakhrushev’s group recently applied data-independent acquisition (DIA) for direct analysis of O-glycoproteins, which was a major advance in characterizing glycosites and glycans(Ye et al, 2019). However, these methods are all based on enrichment of certain types of glycans, mainly GalNAc-type glycans. Therefore, only GalNAc-type O-glycosylation has been heavily studied. A non-biased comprehensive profile of glycan structures and O-glycosites at the whole proteome level has still not been satisfactorily achieved. High-resolution mass spectrometry-based proteomics has become the most powerful tool to study protein glycosylation. Sensitivity, sequencing speed and peak capacity are the key factors for achieving in-depth identification of the glycoproteome. The newly emerging trapped ion mobility spectrometry (TIMS) coupled with TOF mass spectrometry (TIMS-TOF) uses a revolutionary data acquisition technology called Parallel Accumulation - Serial Fragmentation (PASEF) which increases the sampling duty cycle to 100%. This scanning speed, together with the capability to separate isomeric peptides, pushes the instrument sensitivity to a new height and thus makes it possible to analyze the glycoproteome at a much deeper level(Meier et al, 2015).

## Results

### Enrichment-Free Identification of Native Definitive (EnFIND) O-glycoproteome

Taking advantage of the high sensitivity and high isomer separation capability of TIMS-TOF with PASEF, we herein developed an enrichment-free method to identify the native definitive (EnFIND) O-glycoproteome. We applied EnFIND to a whole cell lysate of A549 and human serum. To further challenge the sensitivity of this method, we also applied it to exosomes derived from cells or serum, which can only be purified in very limited amounts.

The workflow of EnFIND is shown in Figure 1A. A small amount (200 ng) of each sample was loaded directly into a nanoLC-TIMS-TOF after a standard pre-treatment process with dual-enzyme digestion to increase the peptide sequence coverage. The general mode of TIMS operation included two steps with ion accumulation (TIMS 1) and serial elution (TIMS 2) of ions separated on the basis of ion mobility from the TIMS device by decreasing the electrical field. The heatmap shows the nested m/z and ion mobility distribution in the PASEF scan, which, together with retention time (RT), provides 3-dimensional (3-D) peptide information for the database search. The 3-D data and the 100% duty cycle of PASEF are the two unique features that bring the sensitivity and thus the protein identification to a new level.

**Figure 1.**
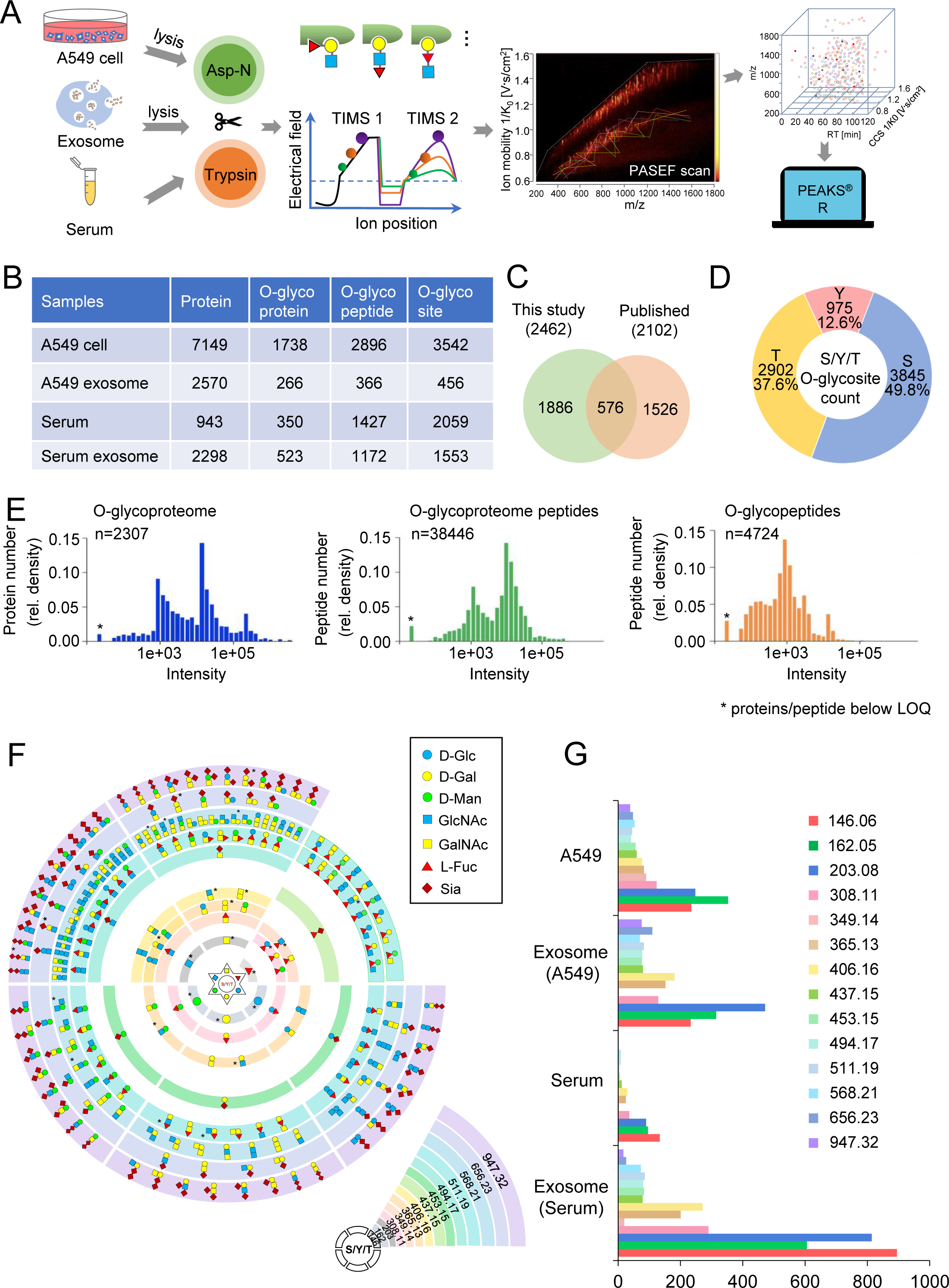
Enrichment-Free Identification of Native Definitive (EnFIND) O-glycoproteome. A. Workflow of EnFIND. B. Summary of the number of identified proteins, O-glycoproteins, O-glycopeptides and O-glycosites. 5% false discovery rate (FDR) at the protein level was used as the cut-off. C. Overlap of the O-glycoproteins identified in this study with previously published O-glycoproteins. D. The relative proportion of serine/tyrosine/threonine (S/Y/T) O-glycosites identified in this study. E. Histograms of the O-glycoproteome, O-glycoproteome peptides, and O-glycopeptides. Y-axis is the percentage of protein or peptides number to total protein or peptides number; X-axis is the intensity. The proteins or peptides with intensity lower than 2e+02 are indicated by an asterisk. F. Systematic illustration of O-glycoforms selected in this study. O-glycoforms reported in the literature are indicated by an asterisk. G. The distribution of 14 types of O-glycoform in individual samples.

We used the total numbers of identified proteins (IDs) as a benchmark for the quality control of EnFIND (Fig 1B). The total 7149 and 943 protein IDs were identified in the A549 whole cell lysate and human serum, respectively. Under these conditions, we identified a total of 1738 and 350 O-glycoproteins in A549 whole cell lysate and human serum, respectively (Fig 1B). In combination of all samples analyzed, we identified a total of 2462 O-glycoproteins compared to the 2102 O-glycoproteins from recently published Glyco-DIA library. Among 2462 O-glycoproteins, 1886 O-glycoproteins specifically in this study (Fig 1C)(Ye et al, 2019). Moreover, 148 and 431 O-glycoproteins were identified exclusively in exosomes derived from A549 or serum (Fig EV1, EV2). This substantially increases the existing knowledge of the O-glycoproteome. Gene ontology (GO) analysis of the identified glycoproteins in cells and serum showed that various localizations, activities and functionalities are associated with O-glycoproteins, consistent with their important role in different aspects of biology (Fig EV3). The distribution of O-glycosites is consistent with previous reports, with the O-linkages occurring more frequently on serine (49.8%) and threonine (37.6%) than on tyrosine (12.6%) (Fig 1D).

The high sensitivity of TIMS-TOF with PASEF is one of the key aspects of the EnFIND. O-glycoproteome and total peptides of O-glycoproteome share the same trend of distribution at 1e3 to 1e5 and 1e3 to 5e4, respectively (Fig 1E, left and middle panels). However, O-glycopeptides show a different distribution pattern, with the relative density peaks shifted to 1e2 to 5e3, and only one peak at intensity 1e3 (Fig 1E, right panel). This demonstrates a successful coverage of low-abundance O-glycopeptides by TIMS-TOF with PASEF. Our data suggest that EnFIND delivers deep coverage of complex native samples.

In the EnFIND approach, we optimized the glycoform database search by constructing 14 types of O-glycoforms derived from the 6 monosaccharides based on previous literatures suggested high structural heterogeneity (Fig 1F)(Joshi et al, 2018; Sheikh et al, 2017). We found that GalNAc/GlcNAc (203.08 m/z) O-glycosylation is indeed a major type, but other types of O-glycosylation are also abundant (Fig 1G).

### Features of the EnFIND method

The ion suppression of very low-abundant O-glycosylated peptides by unmodified peptides is a major roadblock in the identification of O-glycoproteins. In the EnFIND method, we leveraged the capability of separation of isomers by ion mobility spectrometry to achieve full resolution of co-eluted O-glycopeptides and the corresponding unmodified peptide. The extracted ion chromatogram in Figure 2A shows that the O-linked GalNAc/GlcNAc peptide DSKPDTtAPPSSPK (precursor *m/z* 815.895 *z* = +2) and its unmodified form DSKPDTTAPPSSPK (precursor *m/z* 714.35 *z* = +2) cannot be separated by regular reversed-phase chromatography, but they are well distinguished on the basis of ion mobility, presented in extracted ion mobilogram (EIM).

**Figure 2.**
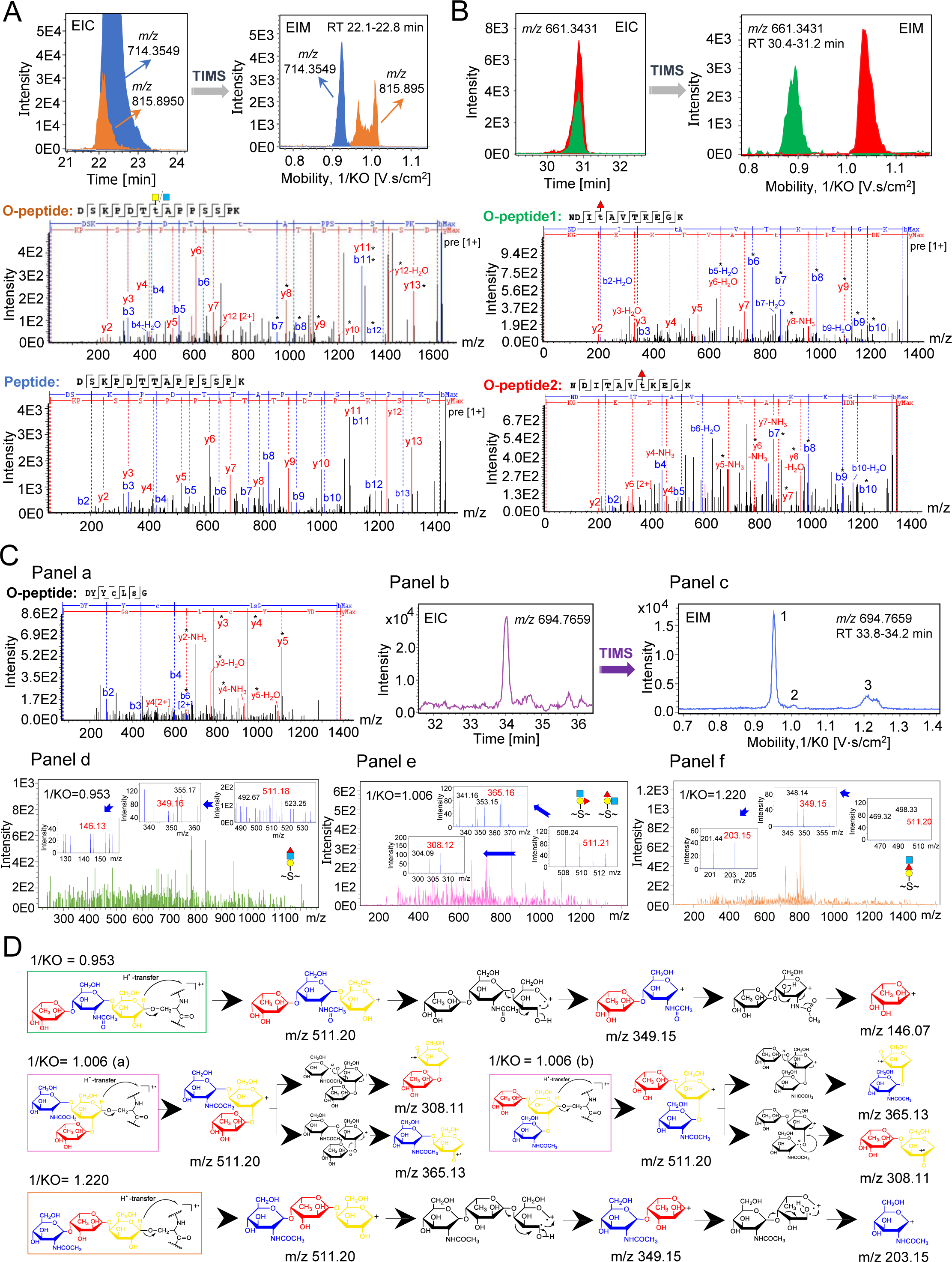
Features of the EnFIND method. A. Ion mobility separation of the O-glycopeptide DSKPDTtAPPSSPK ([M+3M]^3+^, 815.895 m/z) and the unmodified peptide DSKPDTTAPPSSPK ([M+3M]^3+^, 714.355 m/z). The mass of GlcNAc (blue square) or GalNAc (yellow square) is 203.08). Upper panel: Extracted ion chromatogram (EIC) and extracted ion mobilogram (EIM). (blue: DSKPDTTAPPSSPK; orange: DSKPDTtAPPSSPK). Lower panel: MS/MS spectra of DSKPDTtAPPSSPK (O-peptide) and DSKPDTTAPPSSPK (peptide). The characteristic peaks are indicated by an asterisk. B. Ion mobility separation of two isomeric O-glycopeptides, NDItAVTKEGK and NDITAVtKEGK ([M+3M]^3+^, 661.343 m/z). The mass of Fucose is 146.06 (red triangle). Upper panel: EIC and EIM of NDItAVTKEGK (green) and NDITAVtKEGK (red); Lower panel: MS/MS spectra of NDItAVTKEGK (O-peptide1) and NDITAVtKEGK (O-peptide 2). The characteristic peaks are indicated by an asterisk. C. Determination of different glycan structures of isomeric O-linked trisaccharide (mass = 511.19) glycopeptides. Panel a: MS/MS spectrum of DYYCLsG ([M+2M]^2+^, 694.766 m/z) (O-peptide). The characteristic peaks are indicated by an asterisk. Panel b: EIC of DYYCLsG ([M+2M]^2+^, 694.766 m/z) Panel c: EIM of the total separation of DYYCLsG. **1** (Mobility, 1/K0 = 0.953), **2** (Mobility, 1/K0 = 1.006), and **3** (Mobility, 1/K0 = 1.220) indicate different glycan isomers at the site. Panel d: MS/MS spectrum of **1** (CID fragment ions of 349.16 m/z and 146.13 m/z, corresponding to the sequentially loss of a Gal/Glc/Man and GlcNAc/GalNAc residues). Panel e: MS/MS spectrum of **2** (CID fragment ions of 365.13 m/z and 308.11 m/z, corresponding to the loss of Fucose and GlcNAc/GalNAc residues). Panel f: MS/MS spectrum of **3** (CID fragment ions of 349.15 m/z and 203.15 m/z, corresponding to the sequentially loss of a Gal/Glc/Man and Fucose residues). D. Proposed cleavage mechanism maps for the generation of various linkage-diagnostic fragments from the trisaccharide (mass = 511.19) glycopeptides of **1** (1/K0=0.953), **2** (1/K0=1.006) and **3** (1/K0=1.220). The components of the trisaccharide are fucose (red color, mass = 146.07), GlcNAc/GalNAc (blue color, mass = 203.15) and Gal/Glc/Man (yellow color, mass = 162.05)

Identification of isoforms is always a “tough spot” for modern mass spectrometry. Most research groups focus on site-specific glycosylation, assisted by the different specific fragmentation features of modern mass spectrometry techniques, such as collisional induced dissociation (CID), electron transfer dissociation (ETD) and/or electron captured dissociation (ECD)(Darula et al, 2016; Halim et al, 2012; Hoffmann et al, 2016; Levery et al, 2015; Nilsson et al, 2009). In the EnFIND approach, TIMS-TOF with PASEF demonstrates the power of ion mobility spectrometry to identify isomeric O-glycopeptides. We distinguish two types of isomeric peptides in O-glycosylation: adjacent sites modified with the same glycoform and isomeric glycoforms at the same site. The two EIM peaks of DSKPDTtAPPSSPK (precursor *m/z* 815.895 *z* = +2) in Figure 2A indicate two isomeric glycoforms of GlcNAc and GalNAc, with mass of 203.08, added to the same threonine residue (T). Co-eluting isomeric O-linked fucose (mass = 146.06) peptides of NDItAVTKEGK and NDITAVtKEGK show a total separation in ion mobility, resulting in high-quality MS2 spectra that enabled accurate localization of the two different threonine (T) sites (Fig 2B). More excitingly, we are able to identify complex glycan isomers. Figure 2C shows an MS2 spectrum of the doubly charged precursor peptide of DYYCLsG (precursor *m/z* 694.77) with an O-linked trisaccharide glycan (mass = 511.19) modified at residue of serine (S) (Fig 2C, panel a). The EIC shows only one peak for the precursor, however, a full separation of the isomers of this trisaccharide glycan was visible in the EIM (peaks 1, 2, and 3) (Fig 2C panel b, c). This trisaccharide glycan was composed of three monosaccharides, with mass of 146.06, 162.05 and 203.08 respectively. From the characteristic fragments identified in the MS2 spectra, we are able to deduce the most likely glycan structures corresponding to each peak (Fig 2C, panels d-f). The proposed glycan isoforms are listed in Figure 2D. The electron transfer diagrams show the most favorable trajectory of all fragments to each isoform, thus further supporting the glycan structures. Although we still cannot distinguish the monosaccharide Glc, Gal and Man (mass =162.05) or GlcNAc and GalNAc (mass = 203.08) in this trisaccharide glycan, our method achieves the highest resolution in isomer analysis in proteomics studies. The further confirmation of glycan structures will require release of glycans from purified glycoproteins followed by mass spectrum analysis and more information on chemical biology and glycosidic bond dissociation energy(Chai et al, 2018; Tang et al, 2018; Yu et al, 2012).

Antibodies are highly glycosylated proteins. Glycosylation on the variable and constant regions of antibodies is known to influence antigen binding, stability and effector functions, which are related to a variety of diseases(Arnold et al, 2007; Reily et al, 2019). Compared to N-glycosylation, antibody O-glycosylation, especially on the proteomic level, has received less attention due to the lack of detection methods(de Haan et al, 2020). The sensitivity advantage of EnFIND prompted us to investigate the O-glycosylation of antibodies in human serum.

### Identification of O-glycosylation on antibodies

We analyzed the O-glycosylation of antibodies in whole human serum, including total serum, high abundant proteins fraction, exosome fraction, and antibodies purified by protein A/G. Combining the data, we found that antibodies are highly O-glycosylated on all regions (Fig 3A). 110 out of 144 total antibodies are O-glycosylated with 701 glycosites identified on both variable and constant region on heavy and light chain of antibodies with multiple glycoforms (Fig 3B, Table EV1). We identified O-glycosylation of all five isotypes, IgG, IgA, IgM, IgE and IgD (Fig 3C, Table EV2). O-glycosylation of IgD and IgE are rarely reported, consistent with the low amount of IgD and IgE in human serum. The complementarity determining region (CDR), also named hypervariable region, of an antibody determines the antigen binding affinity and specificity. From the IMGT antibody database, we found that the relative frequencies of serine, threonine and tyrosine are high in CDR1 and CDR2 (Fig EV4). We detected 49 antibodies with O-glycosylation in CDR. Furthermore, 100 O-glycosites were identified including 38 isomeric glycopeptides caused by the same modifications on adjacent sites, and 28 glycoform isomers (Fig 3D). The MS2 spectrum of each identified isomeric glycopeptides was manually confirmed (Fig EV5). The heterogeneity of O-glycosylation on CDRs may further add to the diversity of the antibody repertoire.

**Figure 3.**
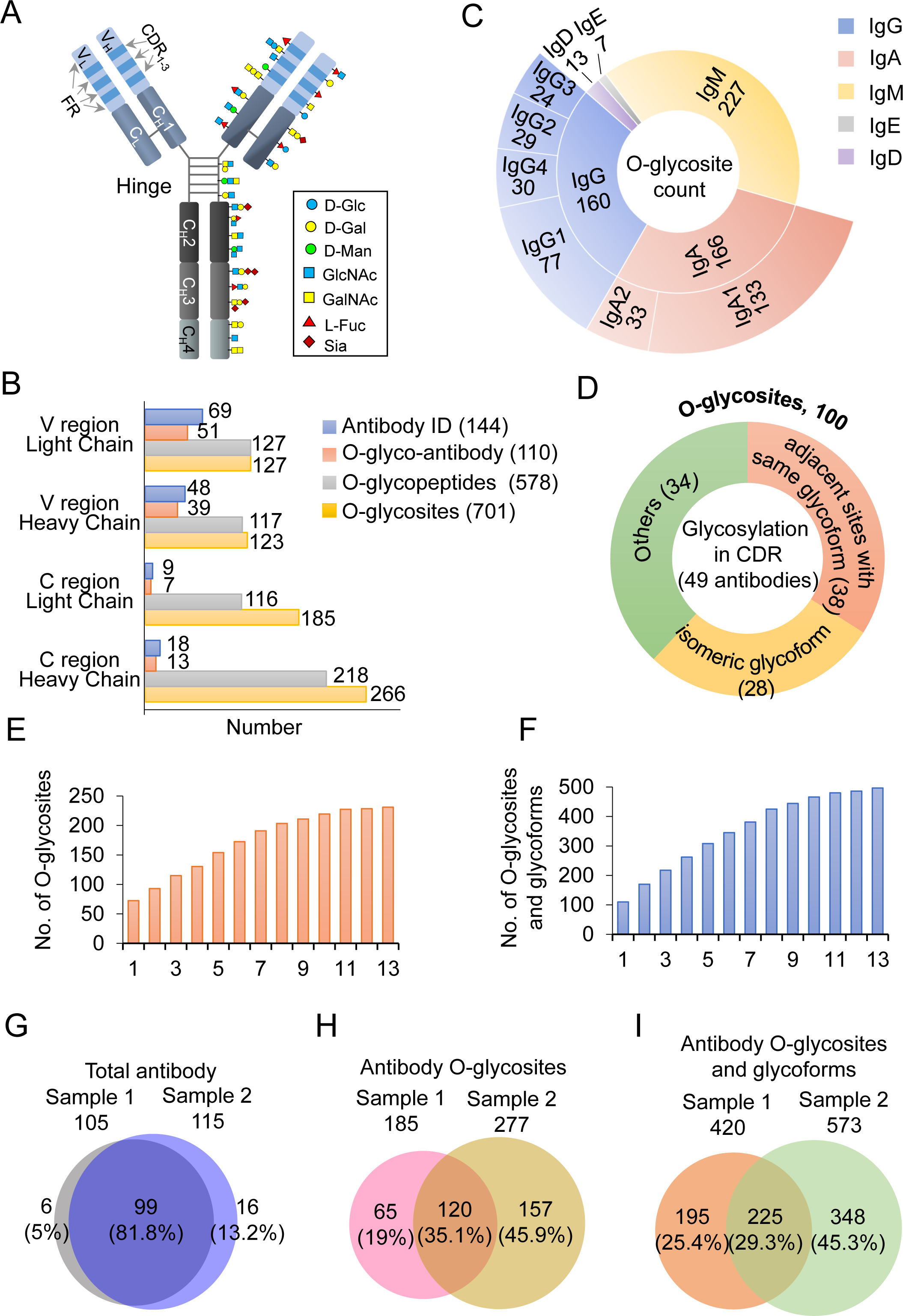
Identification of O-glycosylation on antibodies. A. A schematic overview of identified O-glycosylated antibody from serum and serum exosomes. O-glycosylation sites with multiple glycoforms were identified in the variable domain, constant domain and hinge region. B. Summary of total O-glycosylations of antibody in variable and constant regions. C. Summary of total number of O-glycosylation sites in different categories of antibody in human serum samples. D. Summary of isomeric O-glycopeptides in CDRs. E, F. Average numbers of antibody O-glycosites (E) or O-glycosites and O-glycoforms (F) identified in two healthy controls. Each sample was repeatedly analyzed 13 times. G, H, I. Comparison of identified antibodies (G), antibody O-glycosites (H) and antibody O-glycosites and O-glycoforms (I) in two healthy controls. The data were the union of 13 analyses.

We found that antibody was O-glycosylated at multiple sites and with different glycoforms. Thus, we hypothesized that antibody O-glycosylation raise antibody diversity in addition to amino acid sequence. We build a comprehensive map of antibody O-glycosylation in human serum from two healthy controls using EnFIND approach. The average number of O-glycosites or O-glycosites and O-glycoforms reached a plateau after 13 repeat runs (Fig 3E, F), with an average of 231 and 496 respectively in the two samples tested. What intrigued us is that the total antibodies ID identified in the two healthy controls are quite similar, with 81.8% overlap (Fig 3G). Surprisingly, the O-glycosites and the O-glycosites with different glycoforms overlap by only 35.1% and 29.3%, respectively (Fig 3H, I). These data demonstrated that antibody O-glycosylation in the serum are highly diverse. The significance of this is currently a mystery.

### Antibody O-glycosylation is correlated with autoimmune diseases

Alterations of antibody glycosites or glycoforms can contribute to immune dysregulation and a range of autoimmune and chronic inflammatory diseases. However, current studies focus on antibody N-glycosylation, and the O-glycosylation status is rarely determined(Reily et al, 2019). The EnFIND approach detect the O-glycosylation level of proteins at the native state. We next explored antibody O-glycosylation in diseases, with the aim of identifying disease-associated O-glycosylation patterns. We applied EnFIND to study the autoimmune diseases. Using the workflow described above (Fig 1A), we first selected the constantly identified O-glycosites. Then, we obtained the O-glycosylation level by calculating the ratio of total peptide-spectrum matches (PSM) of O-glycopeptide to the total PSM of the corresponding peptide. We analyzed 10 systemic lupus erythematosus (SLE) serum samples and 10 healthy controls (Table EV3). This allowed us to identify statistically different O-glycosylations on 8 O-glycosites in SLE which are located at hinge region of IGG1 (T225, T227), CH2 region of IGG1 (S241,T252), CH3 region of IGHG2 (S257), and constant regions of light chain IGLC7, IGLC3 and IGKC (Fig 4A, Fig EV6, Table EV4). Interestingly, O-glycosylation on S133 of the IGG1 CH1 region, S59 of IGLC3, and S14 and S20 of IGKC are significantly different between the autoimmune diseases rheumatoid arthritis (RA) and sjogren’s syndrome (SS) (Fig 4C, Table EV4). We also studied plasma samples from patients with mild and critical IgA nephropathy, with the aim of finding clues about the transition from the mild cases to critical cases. Plasma samples were analyzed from 8 mild IgA nephropathy cases, 8 critical cases and 10 healthy controls. We found statistically different O-glycosylations on 8 O-glycosites for IgA nephropathy. They are located on the hinge region of IGG1 (S221, T225, T227, S241) and CH2 region of IGG1 (T252, T262), and the light chain constant region of IGLC3 (Fig 4B, Fig EV5, Table EV4). Glycosylation on the antibody light chain has rarely been reported before, but we consistently identified altered O-glycosylation of the antibody light chain in autoimmune diseases. Altered O-glycosylation in the IgA1 hinge region is reported to be related to IgA nephropathy(de Haan et al, 2020). However, our EnFIND approach did not confidently identify the IgA1 hinge region. This might due to the lack of trypsin and AspN cleavage sites in the IgA1 hinge region. We identified that T361 on IGG1, S46 on IGLC3 and T90 on IGLC7 distinguished the mild cases from the critical cases (Fig 4D, Table EV4). It is worth noting that most studies on O-glycosylation mainly focused on Mucin-type (GalNAc-type) O-glycosylation. We identified disease specific O-glycosylation with a variety of O-glycoforms including O-Fucose and O-Glc/Gal/Man as well as disaccharide 308.11 (m/z), composed of Glc/Gal/Man (162.05 m/z) and Fucose (146.06 m/z). Thus, identification of other types of O-glycosylation other than GalNAc-type will be advantageous for disease-related O-glycosylation studies.

**Figure 4.**
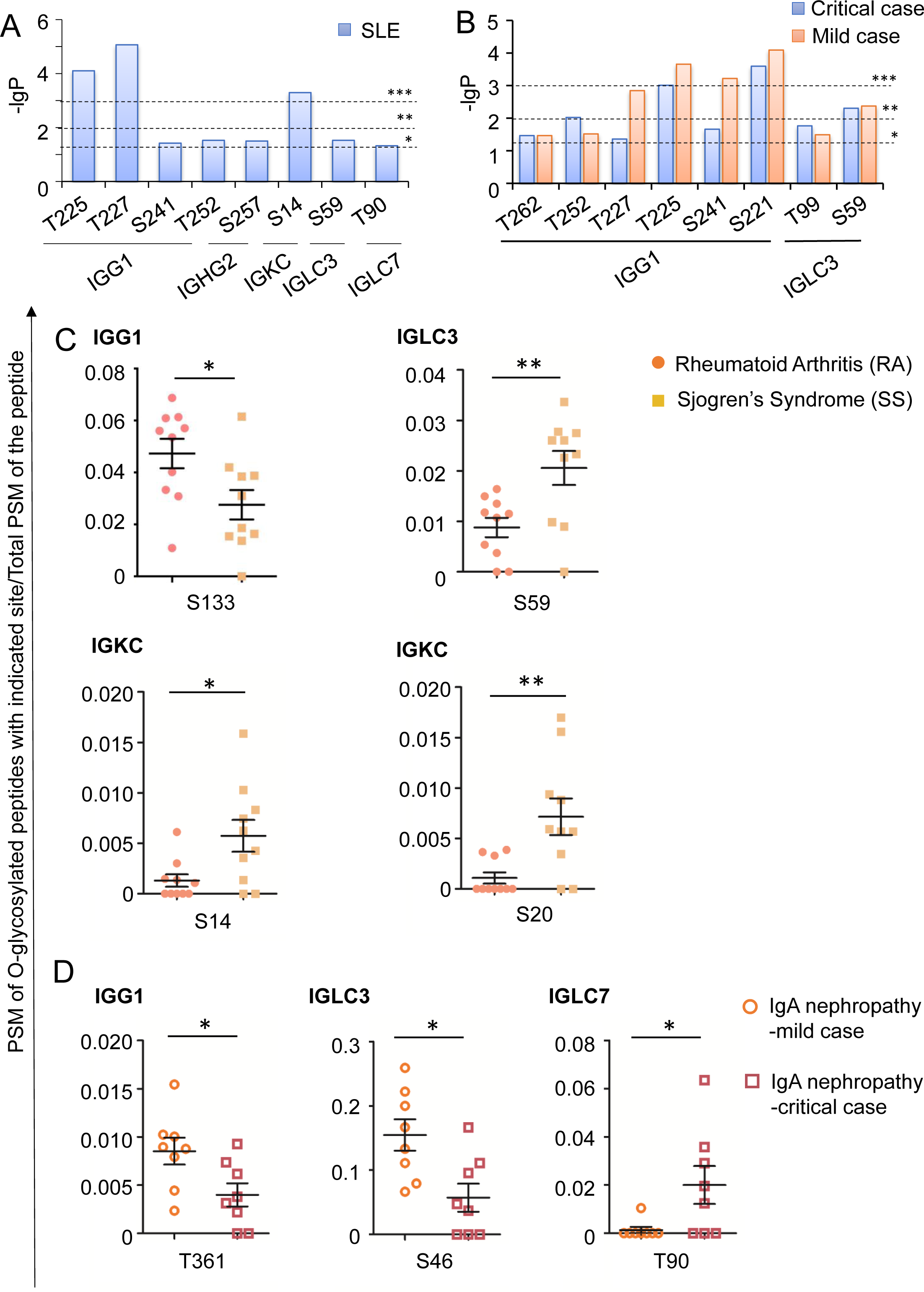
Antibody O-glycosylation is correlated with autoimmune diseases. A, B. Summary of statistically different O-glycosylations in patients with SLE (A) and mild or critical IgA Nephropathy (B) compared to healthy controls by two-tailed, unpaired Student’s t-tests. * P<0.05, ** P<0.01, *** P<0.001. C, D. Scatter plot showing O-glycosites which distinguish RA and SS (C) or mild and critical cases of IgA nephropathy (D). The PSM of O-glycosylated peptide of indicated sites from 3 technical repeats were added and then normalized to total PSM of the indicated peptides added from 3 technical repeats. Individual normalized values for the indicated disease cases and healthy controls are shown. Mean values ± S.E.M, Comparisons were performed by two-tailed, unpaired Student’s t-tests. * P<0.05, **, P<0.01.

## Discussion

We developed an enrichment-free method to identify the native definitive (EnFIND) O-glycoproteome, which enabled us to simultaneously profile the O-glycosites and O-glycoforms from complex native samples. The core advantage of this method relies on the additional separation of isomers on the basis of ion mobility, which greatly decreases the complexity of the sample and further increases the sensitivity. EnFIND requires only trace amounts of sample, making the method suitable for analysis of a wide variety of samples. Without enrichment process, the EnFIND method is detecting the native state, which might reflect the true *in vivo* status of protein O-glycosylation in biological systems, such as whole cell lysates, cell-derived exosomes, serum and exosomes from serum. The 14 types of O-glycoforms included in the data processing greatly increase the number of identified O-glycosites as well as the number of O-glycoforms on one site. Here, we have provided solid evidence that EnFIND is highly effective in separating modified and unmodified peptides and O-glycopeptides isomers, and it can also differentiate isomeric glycoforms. The advantages of EnFIND will guarantee its wide application in O-glycosylation research. Specifically, it will greatly improve O-glyco-library construction for the recently published DIA method for O-glycosylation analysis(Ye et al, 2019).

We applied EnFIND to the analysis of serum samples, and we report for the first time the large-scale O-glycosylation profiling of serum antibodies. We found that antibodies were O-glycosylated in the variable, constant and hinge regions. Most interestingly, we identified a number of O-glycosites located on the CDRs. CDRs play critical roles in epitope recognition and the subsequent immune response, especially CDR3 of the heavy chain, which lies at the center of the antigen-binding site(Vale et al, 2015). Since the sequence of CDR3 is largely missing in database of protein sequence, we only identified a few glycosites on CDR3, but we hypothesize that there should be more O-glycosylation events in this region. So far, we have no evidence of how O-glycosylation on CDRs regulates antibody activity; however, we can speculate that these modifications may regulate antigen binding affinity, antigen-antibody recognition and antibody stability. Different O-glycosites and different types of O-glycoforms, may further increase antibody diversity in addition to primary amino acid sequence. Most notably, antibody O-glycosylation patterns determined a personalized condition among people, the significance of which remains to be determined. The EnFIND method also enables us to investigate the changes in O-glycosylation in diseases. The identified disease-differential O-glycosites provide clues about disease mechanisms. However, the molecular mechanisms underlying the differences in antibody O-glycosylation in autoimmune diseases require extensive studies.

We are now at the initial stage of O-glycoproteome study and there is far more to be explored, especially the functions of O-glycosylation on antibodies. The EnFIND method will benefit biomedical, biopharmaceutical and clinical research. Furthermore, antibodies account for only a small proportion of total O-glycosylated proteins. EnFIND will lead us to a broad identification of protein O-glycosylation in cell, tissue and serum samples to understand how O-glycosylation functions and will boost the knowledge of protein post translational modifications. It will also lead to the discovery of characteristic O-glyco-markers for diseases, which may open up a novel and exciting field for future study.

## Materials and Methods

### Chemicals and reagents

Trichloroacetic acid (TCA), Urea (ACS reagent, 99.0-100.5 %), Iodoacetamide (IAA)and Tris-HCl (ACS reagent) were purchased from Sigma-Aldrich (Merck, USA). RIPAPierceTMlysis, Tris(2-carboxyethyl) phosphine (TCEP), PierceTMtrypsin, AspN protease (MS grade) and BCA Protein Assay Kit were purchased from Thermo Fisher Scientific (San Jose, CA, USA). Formic acid (HPLC grade) was bought from Macklin (Shanghai, China).

### Plasma and serum sample preparation

The participants were recruited from the Peking University People’s Hospital. Serum and plasma samples for research purposes were collected in pro-coagulation or in anticoagulant vacuum tubes using standard venepuncture protocols. Serum or plasma was extracted by centrifugation for 10 min at 3000rpm and subsequently stored at −80 °C before use.

### Protein extraction, reduction, alkylation and digestion

Exosomes were lysed using RIPA buffer containing 1% protease inhibitor at 4 °C for 40 min. After centrifugation at 14, 000 g for 15 min at 4 °C, the supernatant containing exosome proteins was collected. For plasma or serum samples, the high-abundance proteins were immunoisolated by HPLC with a Multi Affinity Removal Column according to the manufacturer’s protocol. The proteins were precipitated with trichloroacetic acid (TCA) and then washed twice with acetone. Proteins were dissolved in 8M urea, then reduced with 10 mM tris-(2-carboxyethyl) phosphine (TCEP) for 40 min and alkylated with 20 mM iodoacetamide (IAA) for 30 min in the dark. Protein digestion was done in two steps. First, the proteolytic enzymes AspN was added in a 1:100 ratio, and incubated at 37 °C for 6 hr. Then, trypsin was added in a 1:50 ratio, and incubated at 37 °C for 16-20 hr. The peptide mixture was desalted on a Monospin C18 column (GL Science, Tokyo, Japan) and dried with a SpeedVac. The dried peptide samples were stored at −80 °C before use.

### LC-TIMS-MSMS analysis

200 ng proteolytic digests were redissolved in 20 μL of 0.1% formic acid (FA) and separated by nanoLC (nanoElute, Bruker Daltonics) with a selfpacked RP column (250 mm x 75 µm, 1.9 µm) at a flow rate of 300 nL/min. The mobile phase consisted of 0.1 % FA in water (A) and 0.1 % FA in acetonitrile (B). The elution gradient used was 2 % to 22 % mobile phase B from 2 to 90 min, 22 % to 37 % mobile phase B from 90 to 100 min, 37 % to 90 % mobile phase B from 100 to 110 min, and then 90% mobile phase B from 110 min to 120 min. The eluates were analyzed online by a trapped ion mobility Q-TOF (timsTOF Pro, Bruker Daltonics) operating in dda-PASEF mode. Mass Range was 100 to 1700 m/z, and 1/k0 was 0.6 V.s/cm2 to 1.6 V.s/cm2. Capillary Voltage was 1700 V to stabilize the electrospray. The dual TIMS analyzer was operated at a locked duty cycle of 100% using equal accumulation and ramp times of 100 ms each. The ramp start voltage Δ6 of TIMS was set to 55 V to avoid possible glycan loss. Each cycle contained one MS1 scan and ten PASEF MSMS scans. Scheduling target intensity was 20000 cts/s, intensity threshold was 2500 cts/s. CID energy started at 20 eV and ended at 59 eV, corresponding to the 1/k0 range of 0.6 V.s/cm2 to 1.6 V.s/cm2. Singly charged precursors were excluded by their position in the m/z-ion mobility plane.

### Data analysis

All raw files were analyzed using PEAKS online software. Experiment type was set as TIMS-DDA with CID fragmentation. The data was searched in the Uniprot human protein databases (140220 entries). The maximum mass deviations of parent and fragment ions after mass recalibration were set to 20 ppm and 0.02 Da, respectively. Trypsin and AspN were chosen as the digestion enzymes, and the maximum missed cleavage was set at 2. Methionine oxidation was set as variable modifications, and Cysteine carbamidomethylation was set as fixed modification. Additionally, the 14 types of O-glycoforms on Ser/Thr/Tyr residues were set as variable modifications. Each valid protein hit should contain at least one unique peptide. The false discovery rates of PSM was set within a threshold value of 0.01. The PSM was manually checked to confirm the O-glycosylation according to b- or y-characteristic fragments containing the O-glycosylated site and the glycan oxonium ions.

### Isolation of exosomes from cultured cells/serum

#### Cultured cells

A549 cells were grown on 150 mm dishes (NEST) in DMEM media with 10% FBS depleted of exosomes until they reached a confluency of 90-100%. One batch of purification required 30 dishes. Conditioned medium was harvested from the cultured cells. All subsequent manipulations were performed at 4°C. Cells and large debris were removed by centrifugation at 4000 g for 20 min followed by 18,000g for 20 min in 50 ml tubes. The supernatant was passed through a 0.2 µm filter and then centrifuged at 100,000 g for 70 min at 4°C. The crude exosome pellet was washed with 60ml PBS, followed by a second step of ultracentrifugation at 100,000 g for 70 min at 4°C to collect the crude exosomes in the pellet.

#### Serum

Blood samples were collected from normal donors for research purposes in 6-ml pro-coagulation vacuum tubes using standard venepuncture protocols. Human serum was withdrawn from the tube for the purification procedures. A total of 50 ml of cell-free serum, which was obtained by combining the serum from 50 healthy controls, was required for one batch purification. Large debris was removed by centrifugation at 1000 g for 10 min followed by 4000 g for 20 min. Crude exosomes were collected by centrifugation at 100,000 g for 70 min, then washed with PBS.

Further exosome purification was performed by Iodixanol-sucrose density gradient centrifugation, using Optiprep (Sigma-Aldrich, D1556) as the density medium. The procedure can be applied to exosome purification from both cultured cells and serum. The gradient was formed by layering 1 mL of 40% (w/v), 1 mL of 20% (w/v), 1 mL of 10% (w/v), and 1 mL of 5% (w/v) iodixanol solutions on top of each other in a 5 mL open top polyallomer tube (Beckman Coulter). Crude exosomes were resuspended in 200 µL PBS and overlaid on top of the gradient. The gradient was centrifuged at 100,000 g and 4°C for 18 h. 10 fractions of 500 µl each were collected from the top and exosomes were mainly enriched in fraction 7.

### Negative staining

Purified exosome pellets were resuspended in 50-100 µl PBS, then a 5 µl sample of each was mixed with the same volume of 2.5% glutaraldehyde (PB, pH 7.4), and fixed for 30 min at room temperature. The sample was spread onto glow discharged Formvar-coated copper mesh grids (Electron Microscopy Sciences, Hatfield) for about 5 min, then washed with water. The sample was then stained with uranyl acetate for 2 min. Excess staining solution was blotted off with filter paper and copper mesh grids were washed with water. Post drying, grids were imaged at 10-100 kV using a transmission electron microscope H-7650.

### Statistics

The quantification of the PSM of O-glycosylation in Figure 4 was done as follows: three technical replicates were done for each sample. To avoid randomness of O-glycosylation detection, the PSM of a certain O-glycopeptide and the total PSM of the corresponding peptides from the 3 technical replicates were added together. The O-glycosylation state of each O-glycopeptide was calculated by normalizing the O-glycopeptide PSM to the total PSM of the corresponding peptides. Statistical analysis was performed in GraphPad Prism. Error bars in the figures represent the mean ± s.e.m. Comparisons were performed by two-tailed, unpaired Student’s t-tests. P < 0.05 was considered statistically significant. We developed software for the calculations mentioned above for all the identified O-glycopeptides.

## Acknowledgement

We thank Dr. Ning Chen of Bruker Daltonics, Inc. Billerica, MA, for her valuable contributions and ongoing support in developing the TIMS-TOF mass spectrometry applications in our laboratory. We thank Dr. Dawei Liu of Crystlab Tech Corp., Beijing, for his constant support with data analysis. We thank Dr. Ying Li of the Cell Facility in the Center of Biomedical Analysis of Tsinghua University for help with electron microscopy. The research was supported by grants from the Fundamental Research Funds for the Central Universities BMU2017YJ003, BMU2018XTZ002 to Catherine CL Wong, the PKU-Baidu Fund 2019BD007 to Catherine CL Wong, the Ministry of Science and Technology of the People’s Republic of China 2018YFA0507102 to Yang Chen and the National Natural Science Foundation of China 91754108 and 31671395 to Yang Chen. Catherine CL Wong thanks the Research Funds from Health@InnoHK Program launched by Innovation Technology Commission of the Hong Kong Special Administrative Region.

## Author contributions

C.CL.W and Y.C designed the experiments. FL.H, H.Z and XJ.H collected clinical samples. X.S, JH.C, WM.T and JT.G conducted the experiments. X.S, WM.T, JH.C, SX.G, FL.H, H.Z, XJ.H, C.CL.W and Y.C analyzed the data. C.CL.W and Y.C wrote the manuscript.

## Expanded view Figures legends

**Fig EV1.**
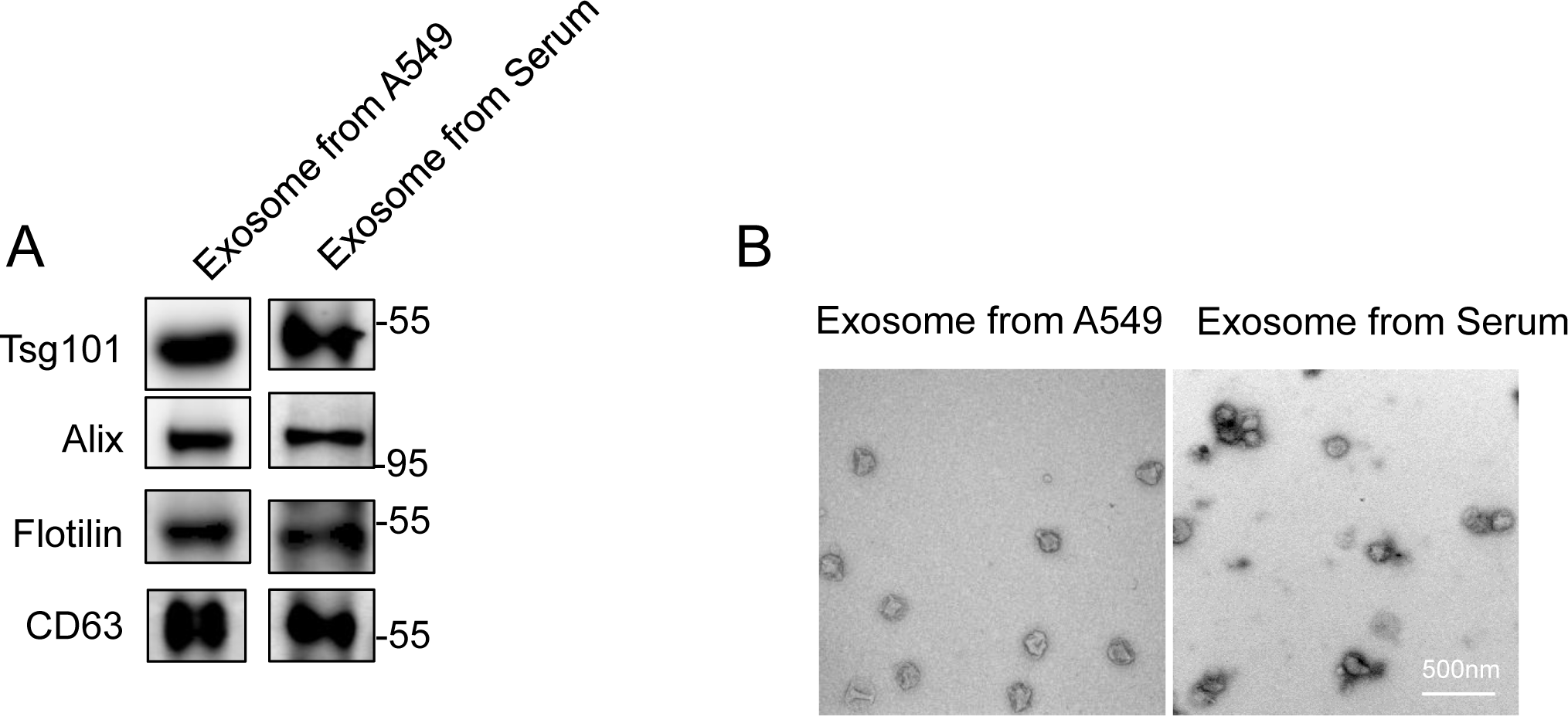
Exosomes purified from cultured A549 cells or serum. (A) Purified exosomes were analyzed by western blotting using antibodies against the exosome specific markers Alix, Tsg101, CD63 and Flotilin. (B) Representative TEM images of negatively stained samples of exosomes. Scale bar, 500 nm.

**Fig EV2.**
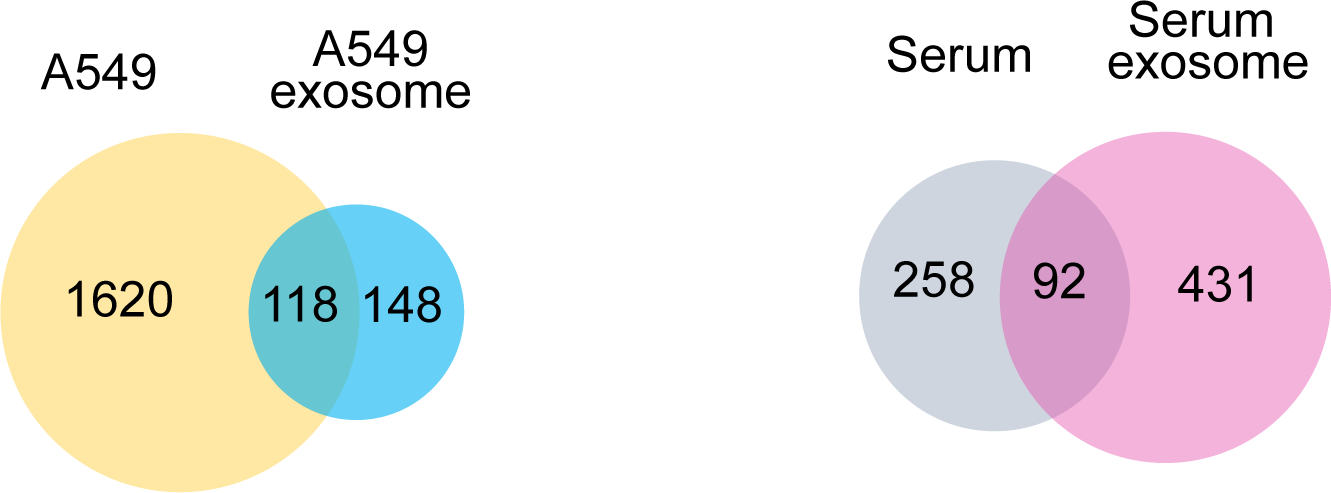
Venn diagram of O-glycoproteins identified in A549 cell lysate and exosomes derived from A549 (left), and serum and exosomes purified from serum (right).

**Fig EV3.**
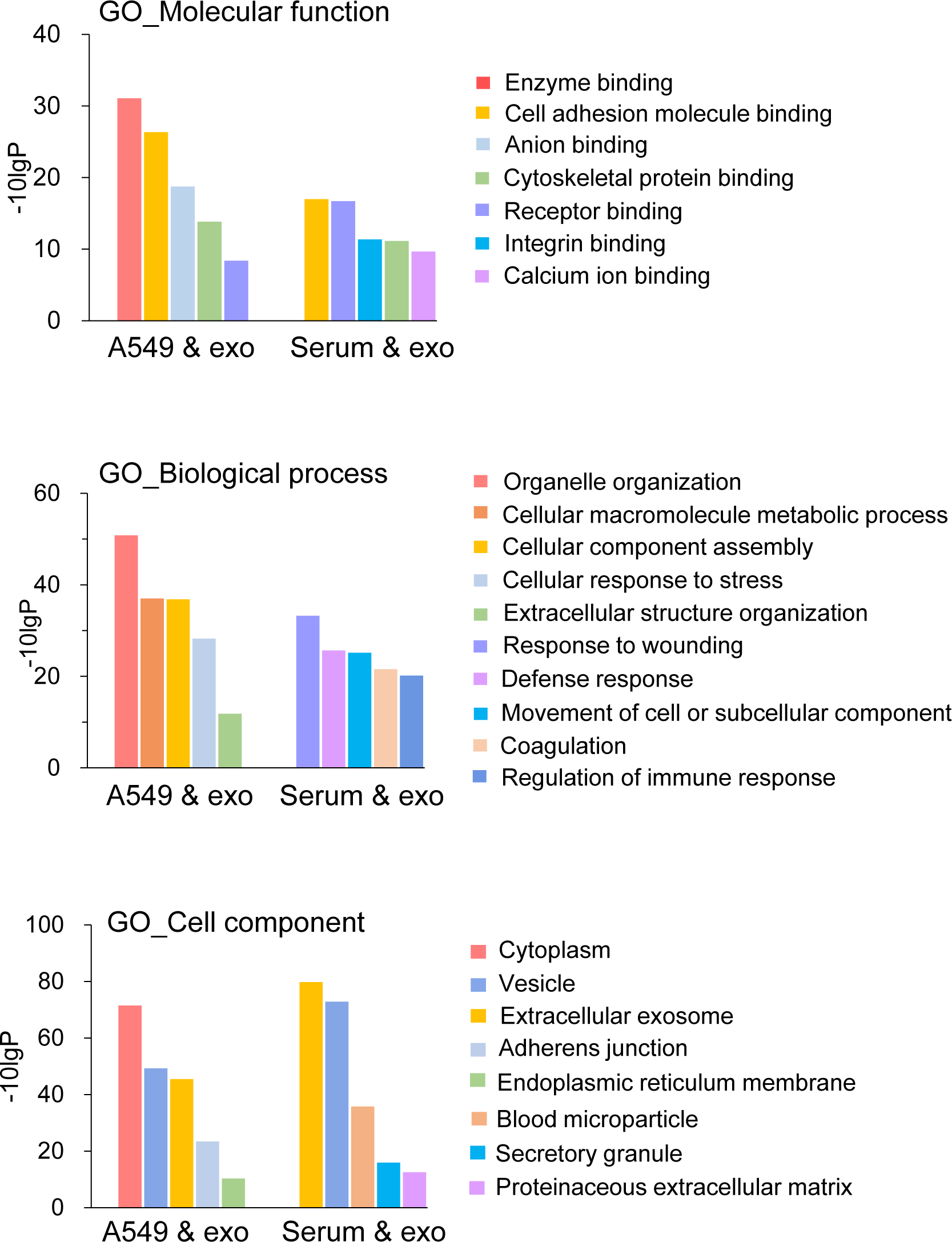
GO analysis of identified O-glycosylated proteins. For GO analysis, the glycoproteins identified in A549 and A549-derived exosomes were combined (A549 & exo) and the glycoproteins identified in serum and exosomes from serum were combined (Serum & exo).

**Fig EV4.**
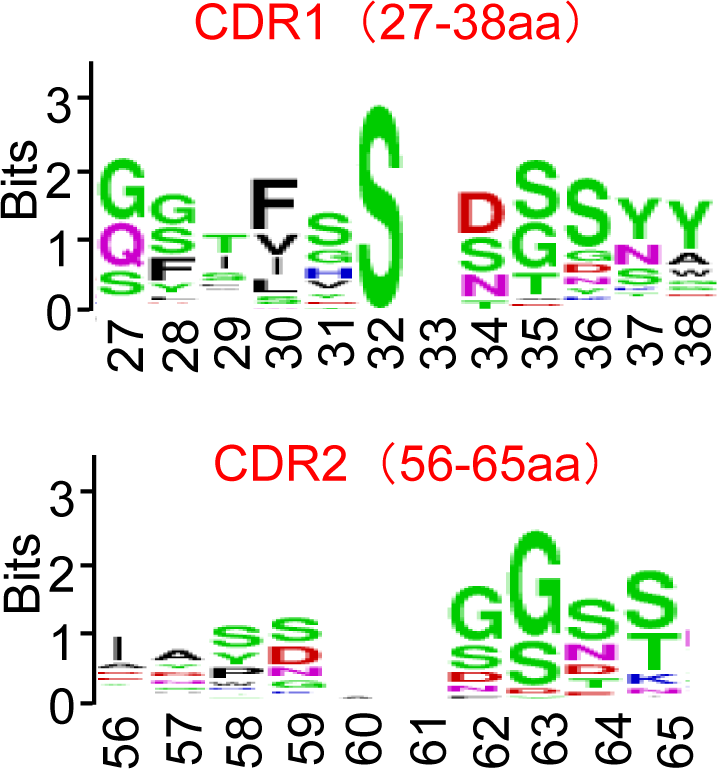
Analysis of amino acid sequences in CDR1 and CDR2 from the IMGT database. The size of the letter represents the frequency of the amino acid.

**Fig EV5 MS2 spectrum of all identified isomeric glycoforms on antibody CDRs.**

**Fig EV6.**
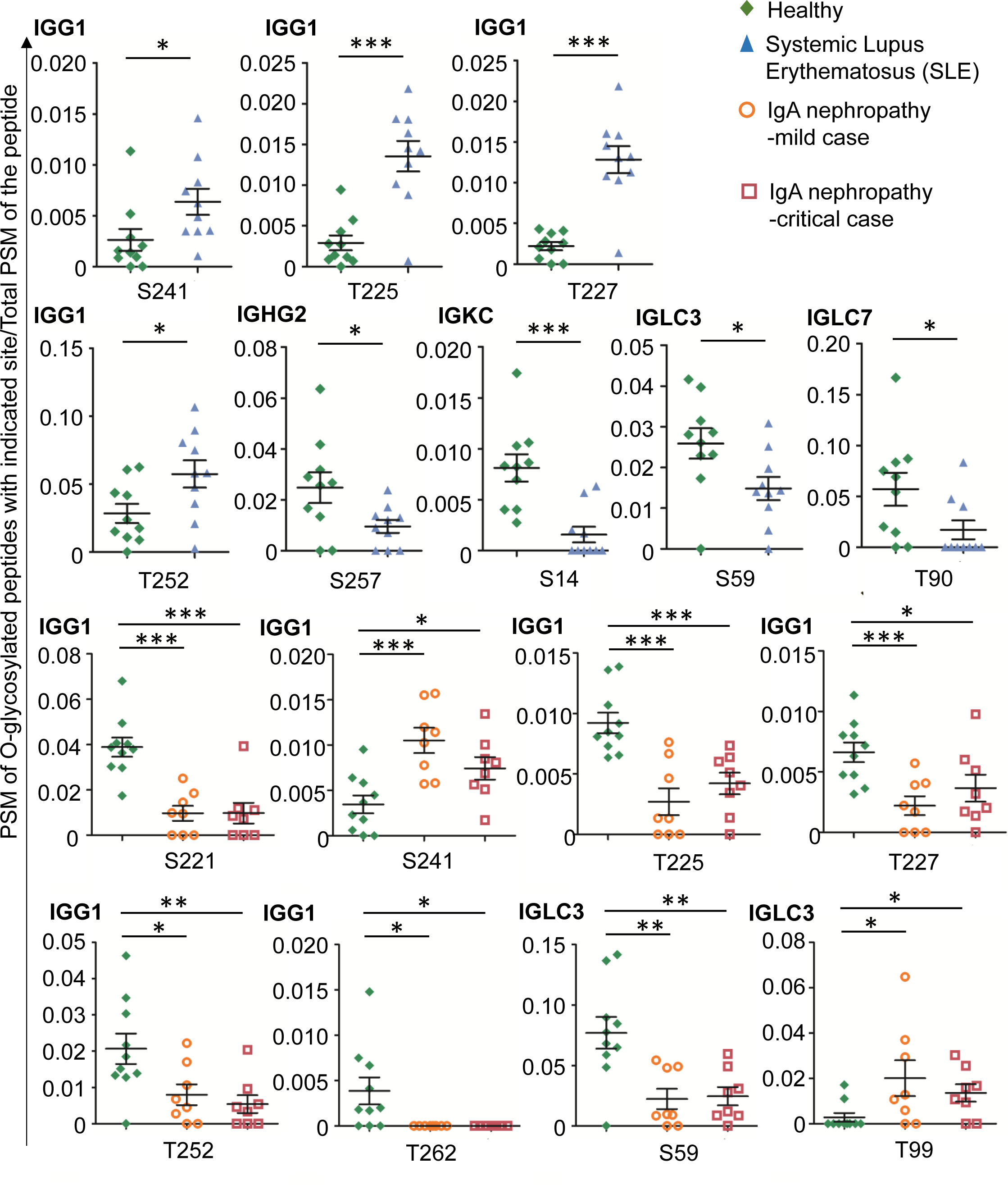
Scatter plot graph of O-glycosites in patients with SLE and mild and critical IgA nephropathy compared to healthy controls. The PSM of O-glycosylated peptide of indicated sites from 3 technical repeats were added and then normalized to total PSM of the indicated peptides added from 3 technical repeats. Individual normalized values for the indicated disease cases and healthy controls are shown. Mean values ± S.E.M, Comparisons were performed by two-tailed, unpaired Student’s t-tests. * P<0.05, **, P<0.01, ***, P<0.001.

## Expanded view table legends

**Table EV1.** Numbers of O-glycoforms in the indicated regions of antibodies in human serum.

| PTM | M.W. | Light chain |  |  |  |  | Heavy chain |  |  |  |  |  |  |  |  |
| --- | --- | --- | --- | --- | --- | --- | --- | --- | --- | --- | --- | --- | --- | --- | --- |
|  |  | Variable region |  |  |  | Constant region | Variable region |  |  |  | Constant region |  |  |  |  |
|  |  | CDR1 | CDR2 | CDR3 | FR |  | CDR1 | CDR2 | CDR3 | FR | CH1 | CH2 | CH3 | CH4 | Hinge |
| Fuc | 146.06 | 11 | 13 | 13 | 23 | 101 | 12 | 3 |  | 59 | 14 | 28 | 11 | 7 |  |
| Man/Hex | 162.05 | 9 | 11 | 10 | 14 | 84 | 2 |  | 2 | 22 | 14 | 42 | 16 | 9 | 11 |
| HexNAc | 203.08 |  | 1 | 6 | 9 | 49 | 2 | 2 |  | 29 | 7 | 39 | 3 | 21 | 14 |
| HexFuc | 308.11 | 1 |  | 2 | 2 | 58 | 6 | 1 |  | 32 | 10 | 2 | 8 |  | 1 |
| HexNAcFuc | 349.14 |  | 1 | 1 | 2 | 19 |  |  |  | 1 |  | 1 |  | 3 |  |
| HexHexNAc | 365.13 |  | 2 | 9 | 6 | 17 |  | 1 | 1 | 1 | 7 | 1 | 6 | 18 | 28 |
| (HexNAc)<sub>2</sub> | 406.16 | 2 | 1 | 10 | 1 | 21 | 2 |  |  | 8 | 8 | 13 | 8 | 14 | 8 |
| NeuAcFuc | 437.15 |  | 4 | 6 | 1 | 44 | 1 |  |  | 23 | 7 |  | 7 | 4 | 5 |
| HexNeuAc | 453.15 |  |  | 2 | 2 | 14 |  |  |  | 11 | 2 | 2 | 9 | 4 | 2 |
| HexNAcNeuAc | 494.17 |  |  | 4 | 1 | 21 | 4 |  |  | 5 | 2 | 1 | 1 | 2 | 3 |
| HexHexNAcFuc | 511.19 |  | 2 | 2 | 6 | 27 | 1 |  |  | 10 | 8 | 3 | 7 | 7 | 7 |
| Hex(HexNAc)<sub>2</sub> | 568.21 |  |  | 6 |  | 11 |  |  |  | 12 | 7 | 3 | 4 | 13 | 36 |
| HexHexNAc(NeuAc)<sub>1</sub> | 656.23 |  |  |  | 1 | 4 |  |  |  |  |  | 3 |  |  | 2 |
| HexHexNAc(NeuAc)<sub>2</sub> | 947.32 |  |  | 4 |  | 6 |  |  | 1 | 1 | 4 | 2 | 6 | 2 | 2 |

**Table EV2.** Numbers of O-glycosites in the indicated regions of different antibody isotypes

| Region | IgG |  |  |  | IgA |  | IgD | IgM | IgE |
| --- | --- | --- | --- | --- | --- | --- | --- | --- | --- |
|  | IgG1 | IgG2 | IgG3 | IgG4 | IgA1 | IgA2 |  |  |  |
| FR | 2 |  |  |  |  | 18 | 4 | 3 |  |
| CDR1 | 9 |  |  |  |  |  |  |  |  |
| CDR3 |  |  |  |  |  |  | 3 | 1 |  |
| CH1 | 8 | 3 | 1 | 4 | 3 |  | 3 | 66 |  |
| CH2 | 35 | 13 | 14 | 15 | 1 | 11 | 1 | 41 |  |
| CH3 | 18 | 13 | 6 | 11 | 19 | 4 | 1 | 15 | 7 |
| CH4 |  |  |  |  |  |  |  | 101 |  |
| Hinge | 5 |  | 3 |  | 110 |  | 1 |  |  |
| O-glycosites | 77 | 29 | 24 | 30 | 133 | 33 | 13 | 227 | 7 |
| O-glycopeptides | 58 | 23 | 20 | 20 | 71 | 28 | 10 | 145 | 5 |

**Table EV3**

Characteristics of the disease and control cohorts.

**Table EV4**

Raw data of the O-glycosylations including total PSM, total O-glycopeptide PSM of the O-glycopeptides mentioned in the study.

